# A G-Quadruplex Containing First Intron Harbors a Functional Polyadenylation Signal in *TP53*

**DOI:** 10.64898/2026.11.08.764788

**Authors:** İrem Cemile Eroğlu, İrem Göller, Elanur Almeriç Yavuz, Özgül Persil Çetinkol, Ayşe Elif Erson-Bensan

## Abstract

First introns of mammalian genes are often longer than other introns and may contain regulatory elements, suggesting that they may have roles in gene regulation. We detected a previously unannotated, low abundance intronic transcript, TP53-INT1, using long-read direct RNA sequencing data. TP53-INT1 has low coding potential and terminates within the first and longest intron of *TP53*. The transcript structure and 3’-end were validated by independent experimental and *in silico* evidence. We identified a canonical polyadenylation signal upstream of the transcript termination site and demonstrated its functionality using reporter assays. Furthermore, a potential G-quadruplex (G4)-forming sequence was identified upstream of this polyadenylation signal. The G4-forming potential was confirmed using spectroscopic and thermal stability analyses. Complementing the *in vitro* characterizations, treatment with Pyridostatin, a G4-stabilizing drug, was associated with decreased *TP53* expression. Together, these findings characterize a previously unannotated *TP53* intronic transcript and reveal a local genomic context combining accessible chromatin, a functional polyadenylation signal, and a potential G4-forming sequence that may influence full-length transcription.

## Introduction

Since its initial identification as a binding partner of the SV40 large T antigen, extensive research has revealed the remarkable functional diversity of TP53 in cancer and normal cells. The evolutionary conservation of *TP53* across even lower eukaryotes further underscores the fundamental importance of its functions [1]. TP53 encodes a transcription factor that responds rapidly to diverse cellular stresses, including DNA damage, ribosomal stress, senescence, telomere erosion, hypoxia, starvation, and oncogene addiction. In unstressed cells, TP53 protein has a short half-life, which is regulated by MDM2 (Mouse Double Minute 2 Homolog) ubiquitin ligase. In response to various stress signals, MDM2 is degraded or inactivated, leading to the accumulation of TP53. Stabilized TP53 forms functional tetramers and acts through both protein-protein interactions in rapid response pathways and transcriptional regulation of a broad range of protein-coding and non-coding genes [2, 3]. Consequently, TP53 orchestrates the stress responses through DNA damage repair, cell cycle arrest, apoptosis, or senescence, thereby earning its well-established title as the “guardian of the genome” [4].

Regulation of TP53 at the protein level is one of the most extensively investigated aspects of TP53 biology, revealing diverse post-translational modifications that contribute to its regulation and activity in normal and cancer cells. In addition, somatic or germline genetic alterations in the TP53 gene, including loss of one or both copies, deletions, many hot-spot missense mutations, and single-nucleotide variants, have been detected in various cancers. Although more than half of all cancers harbor TP53 mutations [4], many tumors retain two intact TP53 copies, indicating other mechanisms may contribute to TP53 inactivation. It is precisely in this context that regulatory elements embedded within the TP53 gene sequence may represent an important, yet relatively unexplored, layer of regulation. In particular, how intronic sequence features may influence *TP53* mRNA levels remains an intriguing area of investigation.

Here, we describe a low-abundance 2.2 kb intronic transcript arising from the relatively long 10.7 kb first intron of *TP53* and terminating after a canonical and functional intronic poly(A) signal. We further identify and validate a G-quadruplex (G4) structure proximal to this poly(A) signal and provide evidence that this structure may modulate *TP53* transcription in cells. Notably, this intronic transcript region identified in this study overlaps with a genomic region previously reported to contain translocation breakpoints in osteosarcomas and patients with Li-Fraumeni syndrome and proposed as a hotspot for genomic rearrangements [5]. Together, these findings highlight an intronic region within the *TP53* locus with potential functional relevance.

## Results

An earlier Oxford Nanopore Technologies (ONT) Direct RNA Sequencing (DRS) analysis of transcriptome complexity in breast cancer cells (PRJNA1195033, PRJNA1196847) [6] identified a previously unrecognized, low-abundance transcript arising from the first intron of *TP53* (Figure 1A). This intronic transcript originates as the full-length *TP53* mRNA but terminates in the first intron (hg38, chr17: 7685249-7687481; hg19, chr17: 7588567-7590799) (Figure S1A). This transcript was not annotated in the Genotype-Tissue Expression (GTEx) database, where *TP53* expression in mammary tissue is shown (Figure 1B). We therefore sought to characterize this transcript; hereafter referred to as TP53-INT1. Additional independent evidence supporting the presence of TP53-INT1 was found in the ENCODE4 Long Read Transcripts and the Human PAS (max version) database, which identified transcript and polyadenylation features consistent with our DRS-based detection of TP53-INT1 (Figure 1C).

**Figure 1.**
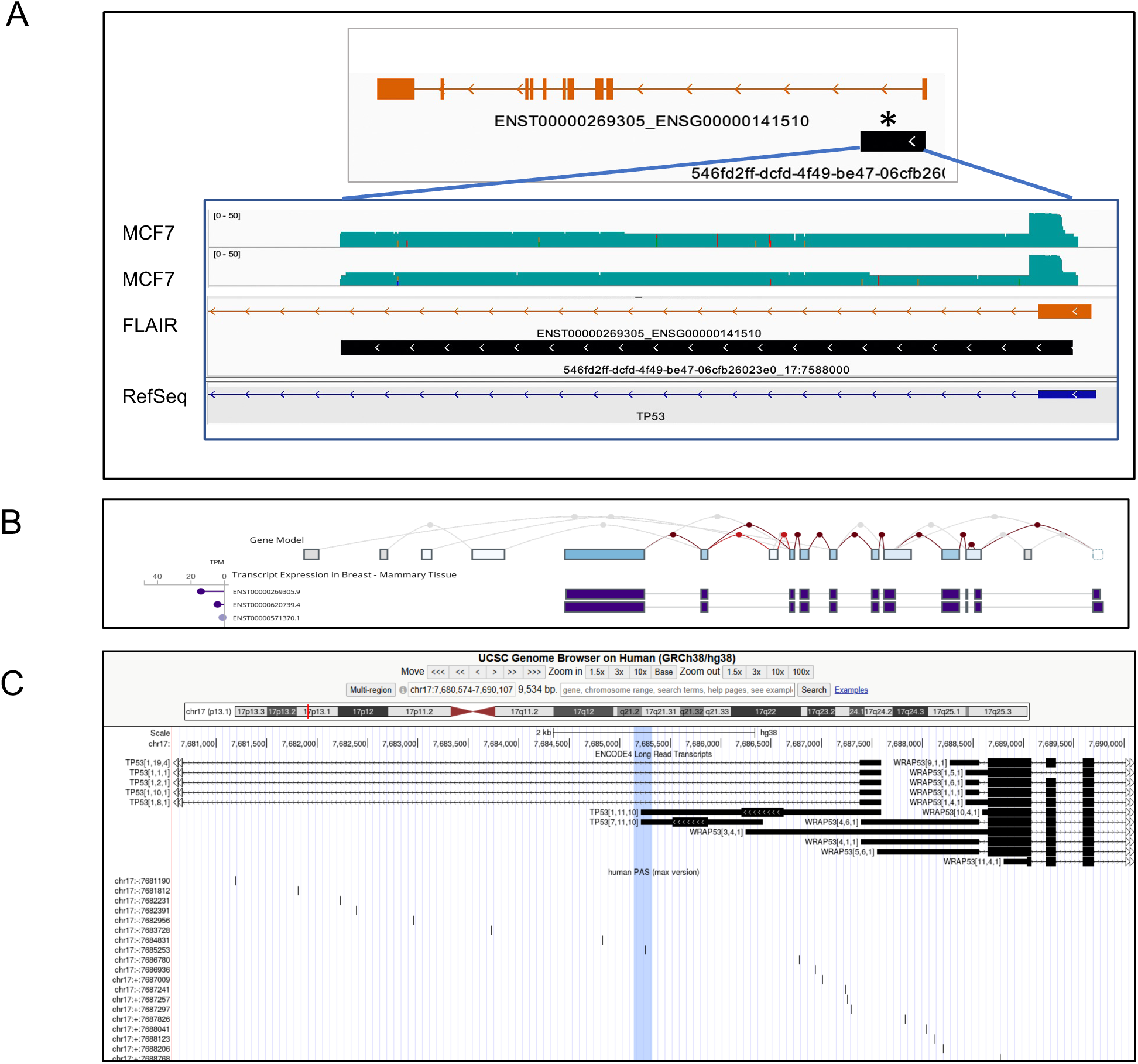
TP53-INT1. A. FLAIR (Full-Length Alternative Isoform analysis of RNA), an algorithm used for long-read RNA sequencing and isoform discovery, identified the annotated *TP53* mRNA (drawn in orange) and an intronically terminated transcript in *TP53* with an unannotated identifier (drawn in black, *). The RNA reads are shown in reference to the RefSeq gene structure of *TP53* in IGV. B. The GTEx database shows annotated *TP53* mRNA isoforms expressed in normal mammary tissue. C. ENCODE4 Long Read Transcripts and Human PAS (max version is an updated version of PolyA_DB (v4) based on 3’READS). The highlighted region shows the 3’-end of TP53-INT1. Please note that TP53 is transcribed from the minus strand.

As TP53-INT1 was identified by native poly(A)+ DRS, it was expected to carry a poly(A) tail. We therefore performed 3’-RACE to validate its 3’-end using an oligo-dT adapter primer during cDNA synthesis and PCR primers for TP53-INT1 and the adapter sequence. Cloning and sequencing of the 3’-RACE PCR product confirmed the DRS-detected, and polyadenylated 3’-end of TP53-INT1 (Figure S1B-D).

Next, using specific primer sets, we performed RT-qPCR to detect TP53-INT1 and *TP53* mRNA levels in a set of breast cancer cell lines (Figure 2A). Because TP53-INT1 resides within the first intron of TP53, we treated all RNA samples with DNase to eliminate genomic DNA contamination and continued with the treatment until no PCR amplification was detected in the absence of reverse transcription (Figure S2). Based on RT-qPCR, TP53-INT1 was detected in all tested breast cancer cell lines. Because this analysis was based on relative expression levels, rather than a direct comparison of the absolute expression levels of the two transcripts, we evaluated the ratio of the two isoforms to assess their relative abundance. Notably, while the TP53-INT1/TP53 relative ratio was in the 2-3 range for most cell lines, the ratio was markedly higher in MDA-MB-157 cells. Of note, MDA-MB-157 cells are known not to express TP53 protein due to a deletion in exon 4 [7].

**Figure 2.**
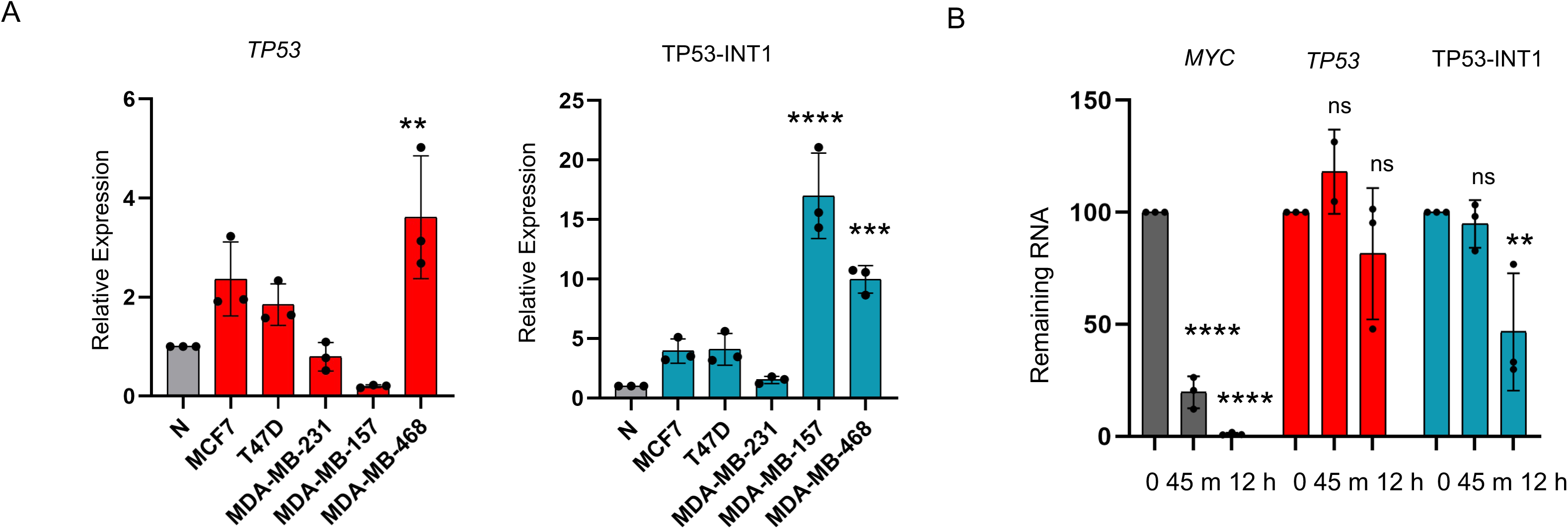
Relative transcript levels. A. TP53-INT1 and *TP53* mRNA levels in breast cancer cell line models detected by RT-qPCR, compared to normal breast tissue cDNA (N). B. RNA stability of *TP53* and TP53-INT1 was detected after actinomycin D (10 µg/ml) treatment in MCF7 cells. RT-qPCR results show the remaining mRNA levels (normalized to *RPLP0*) at 45 min and 12 h of actinomycin D treatment. All time points were compared to the t=0 sample. *MYC* mRNA has a known short half-life and was used as a positive control for treatment. Data represent the mean (SD) of three independent experiments. One-way ANOVA with Tukey’s multiple comparison test was used; ns: not significant, **p<0.01, ***p<0.001, ****p<0.0001.

After confirming TP53-INT1 expression, we assessed its coding potential to determine whether it could encode a protein. Although TP53-INT1 has the potential to encode a 140-amino-acid protein, CPC2 (Coding Potential Calculator 2) classified TP53-INT1 as a non-coding RNA with low coding potential [8] (Figure S3). Despite its low coding potential, TP53-INT1 transcript was relatively stable, as assessed by inhibiting transcription with actinomycin D and measuring the remaining RNA levels by RT-qPCR (Figure 2B). *MYC* mRNA has a significantly short half-life and was used as a positive control in this experiment [9].

To further investigate whether transcription terminates in a manner that could generate TP53-INT1 and account for its detection by DRS, we looked into potential poly(A) signals that would lead to transcriptional termination within the first intron. Investigating the region proximal to the 3’-end of TP53-INT1, we detected two potential poly(A) signals, AAUAAA and UAUAAA, located 30 and 18 nucleotides upstream of the 3’-end of TP53-INT1, respectively (Figure S4). Notably, AAUAAA is the most frequent (canonical) motif and is therefore considered the strongest poly(A) signal in mammalian 3’UTRs, including *TP53* [10, 11, 12]. Poly(A) signals such as AAUAAA are generally located 10-35 nucleotides upstream of mRNA 3’-cleavage sites known as the poly(A) sites (reviewed in [13, 14]). Notably, the 3’-end of TP53-INT1 detected by DRS was a CA dinucleotide, one of the most frequent dinucleotides immediately upstream of the cleavage sites (poly(A) sites) in mammals [15].

To determine the functionality of the intronic poly(A) signals (AAUAAA and UAUAAA) found in TP53-INT1 relative to the TP53 poly(A) signal (AAUAAA) in the terminal exon, we used a *Firefly* luciferase reporter system (pMIR) in which the SV40 poly(A) signal region (50 bp) was removed, generating pMIR-ΔpA. We PCR-amplified and cloned 150-200 bp regions from TP53 intron 1 or the last exon of TP53 into the 3′ UTR of the luciferase reporter in pMIR-ΔpA. The polyadenylation activity of the cloned regions was then evaluated and compared with each other and to that of the SV40 poly(A) signal in the parental pMIR vector. MCF7 cells were also co-transfected with phRL-TK (*Renilla* luciferase) for transfection efficiency normalization, and dual luciferase activities were measured (Figure 3A). Removal of SV40 poly(A) signal region led to a 50% reduction in the *Firefly* luciferase levels in pMIR-ΔpA-transfected cells compared to pMIR-transfected cells. Both of the poly(A) signals from TP53-INT1 and TP53 produced approximately twofold higher luciferase activity than pMIR-ΔpA, reaching levels similar to those observed with pMIR. These results suggested that the cloned sequences rescued the lack of the SV40 poly(A) signal region in pMIR-ΔpA. However, the TP53-INT1 construct had two potential polyadenylation signals, the canonical AAUAAA motif and a downstream UAUAAA sequence. To assess the contribution of the AAUAAA signal, we generated a deletion construct by site-directed mutagenesis so that only the AAUAAA (TP53-INT1ΔpA) motif was deleted. Following co-transfections and a dual luciferase assay, deletion of the AAUAAA signal reduced luciferase activity to a level similar to that of pMIR-ΔpA, suggesting that the intronic AAUAAA motif is the major contributor to the polyadenylation activity for TP53-INT1 (Figure 3B).

**Figure 3.**
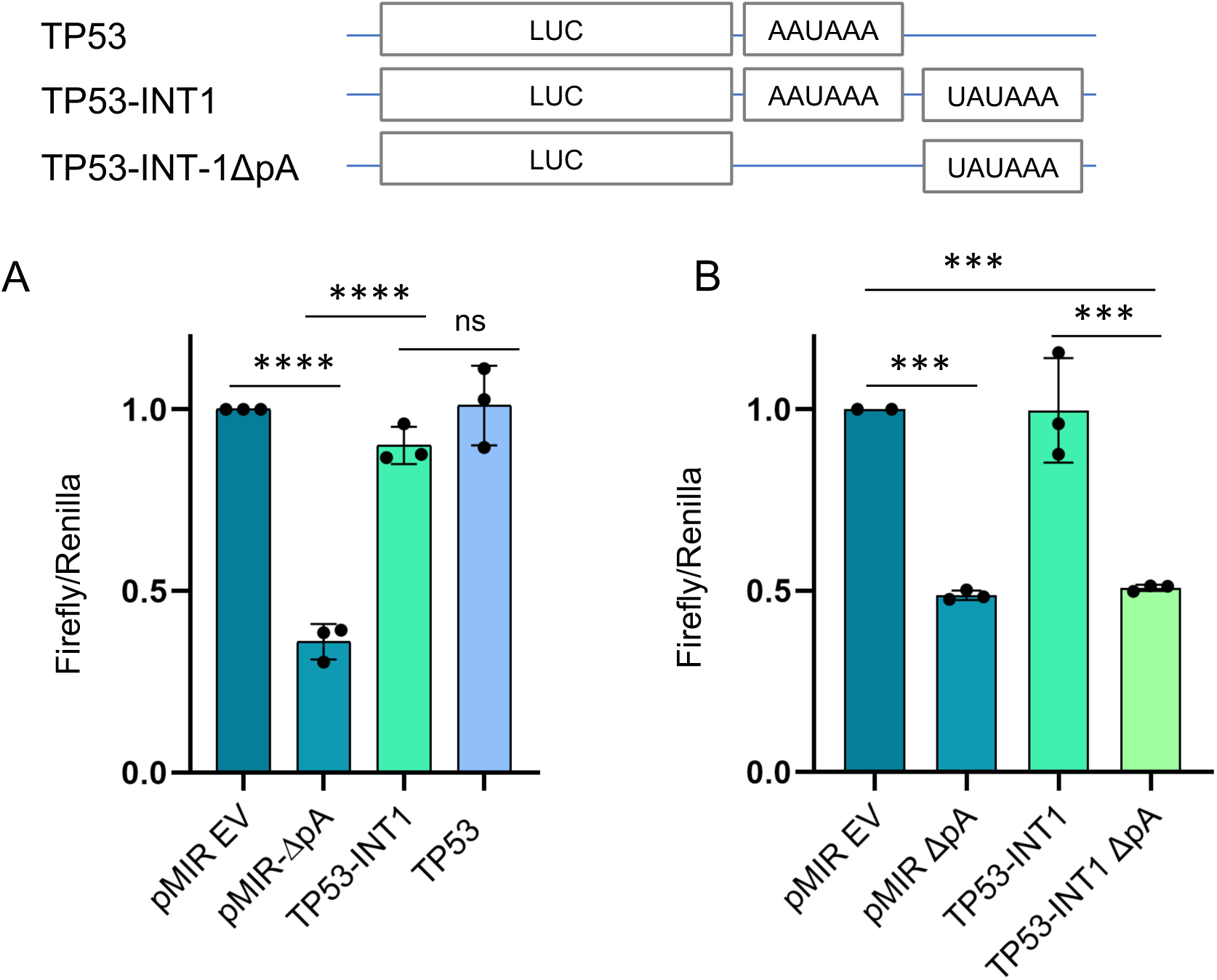
Poly(A) signal activity. The top panel shows the poly(A) signals of TP53 and TP53-INT1 cloned downstream of the *Firefly* luciferase (LUC) gene of the pMIR-ΔpA construct. A. Dual luciferase reporter assay with poly(A) signal regions of *TP53* and TP53-INT1. pMIR has the SV40 poly(A) region, whereas pMIR-ΔpA lacks this region. We transiently transfected MCF7 cells and normalized *Firefly/Renilla* luciferase readouts to pMIR (EV)-transfected cells. B. An independent dual luciferase assay with TP53-INT1 and a deletion mutant that lacks AAUAAA (TP53-INT1-ΔpA) in pMIR-ΔpA. (***p< 0.001, ****p<0.0001, ns: not significant, n=3 independent transfections, one-way ANOVA, Tukey’s HSD)

Next, we sought to investigate how accessible this early intronic poly(A) signal might be to the poly(A) machinery and looked into chromatin architecture and RNA polymerase II (Pol II) occupancy, using publicly available datasets in the Cistrome Data Browser [16]. In particular, Pol II occupancy decreased at the genomic region corresponding to the 3’-end of the TP53-INT1 transcript (Figure 4), accompanied by similar patterns in histone modifications, including H3K27ac and H3K4me3 signals that imply open chromatin status [17]. Given this open chromatin environment, we also examined the sequence upstream of the poly(A) signal region for features that could support non-canonical RNA structures involved in transcription or RNA processing. G4Hunter, a tool for G-quadruplex (G4) prediction, revealed a relatively G-rich sequence 273 nucleotides upstream of the poly(A) signal, prompting us to investigate its potential to form a G4 structure [18]. G4Catchall similarly predicted a G4 structure in this region (Figure S5) [19]. Based on these results, we wanted to experimentally test whether this region proximal to the poly(A) signal in TP53-INT1 has the potential to form a G4 structure.

**Figure 4.**
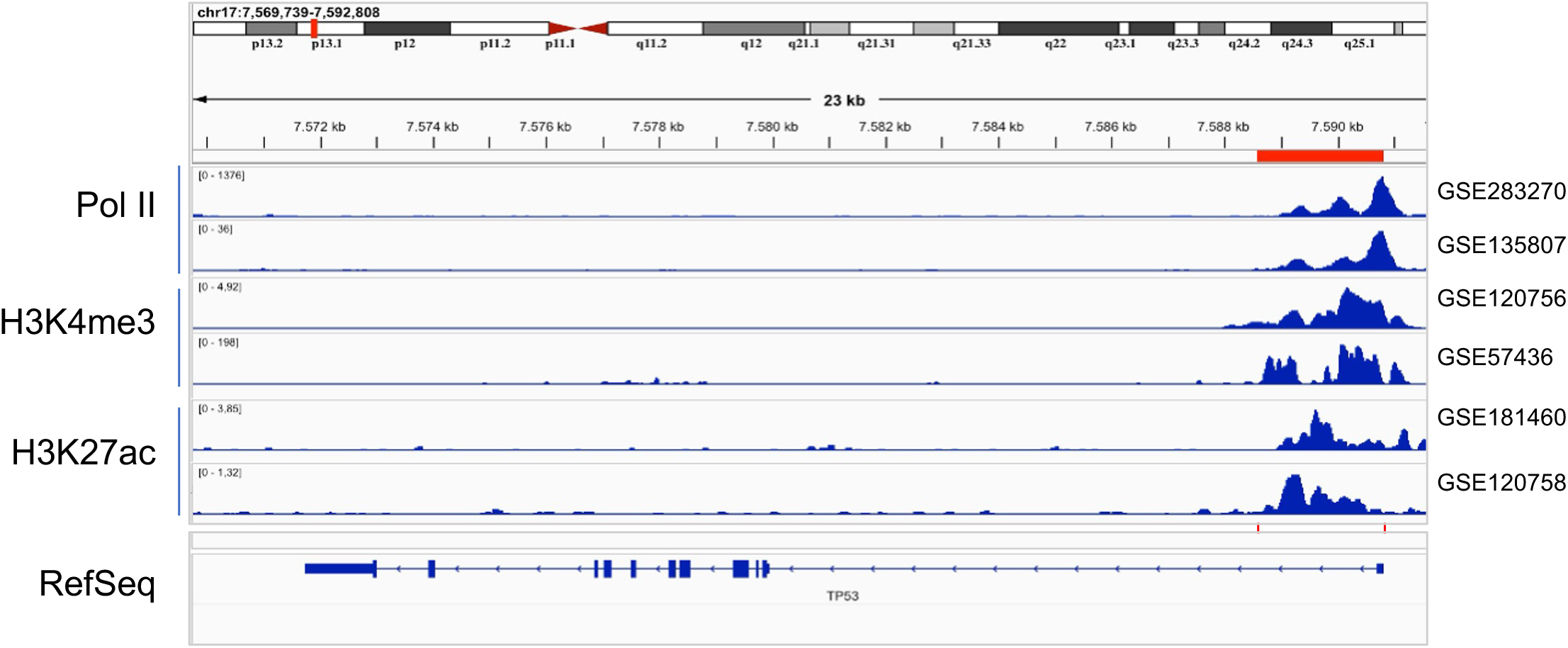
IGV genome browser tracks of Pol II, H3K4me3 and H3K27ac ChIP data from Cistrome datasets for MCF7 (GSE283270, GSE135807, GSE120756, GSE57436, GSE181460, GSE120758). The region highlighted in red shows TP53-INT1.

First, the Thioflavin T (ThT) assay was used to investigate the formation of a potential G4 structure. ThT exhibits selective test fluorescence enhancement upon G4 binding and is therefore commonly used to confirm the formation of G4 [20, 21]. ThT fluorescence emission was measured following incubation with the TP53-INT1 sequence to assess G4 formation. We also tested the complementary strand (TP53-INT1comp), the mutant version of TP53-INT1 (TP53-INT1mut) where guanines were replaced with adenines, the well-characterized G4-forming Pu27 sequence from the *MYC* promoter [22], and a dA_32_ oligonucleotide as a non-G4-forming control (Figure 5A). Incubation of ThT with the putative G4-forming TP53-INT1 sequence resulted in a significant increase in fluorescence in both the presence and absence of KCl. A substantial increase in ThT fluorescence was also observed for Pu27, whereas the TP53-INT1 complementary and mutated sequences, and the dA□□ control oligo displayed low fluorescence at 488 nm, close to the baseline observed for free ThT (Figure 5B). These results were consistent with the formation of a G4 structure within the predicted region of TP53-INT1. For the double-stranded ds (TP53-INT1) sequence, a moderate fluorescence signal was observed. Although ThT is known to bind preferentially to G4 structures, previous studies have also reported a similar weak fluorescence response in the presence of dsDNA [20, 23].

**Figure 5.**
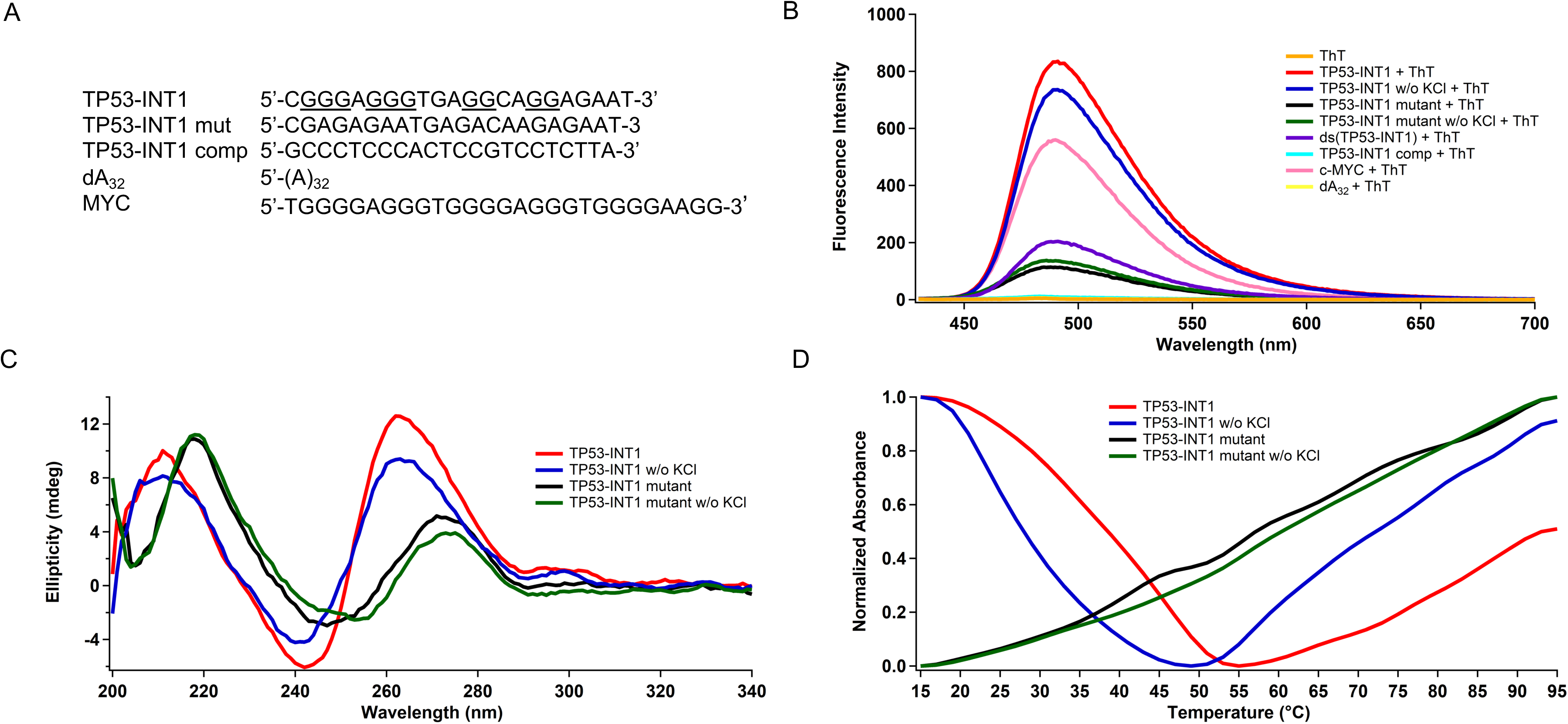
G4 characterization. A. Sequences of oligonucleotides are given. TP53-INT1 (possible G4 forming nucleotides are underlined), TP53-INT1 mut (G to A changes), TP53-INT1 comp (complementary oligo), negative control (d(A)_32_) and positive control (*MYC*, Pu27) oligos are shown, B. Fluorescence spectra of ThT in the presence or absence of oligonucleotides at pH 7.0 in 25 mM K-phosphate buffer with or without additional KCl (pH=7.0), C. CD spectra of TP53-INT1 and TP53-INT1 mutant sequence at pH 7.0 in 25 mM K-phosphate buffer in the presence or absence of additional 70 mM KCl, D. Thermal denaturation profiles of TP53-INT1 and TP53-INT1 mutant oligos at pH 7.0 in 25 mM K-phosphate buffer in the presence or absence of additional 70 mM KCl. Thermal denaturation profile of oligonucleotides obtained from the change in UV-Vis absorbance at 295 nm between 15°C to 95°C.

To further assess the G4 formation, Circular Dichroism (CD) spectroscopy was performed. The three major G4 topologies can be distinguished by their characteristic CD spectra. Parallel G4s typically display a positive band around 264 nm and a negative band around 245 nm, whereas antiparallel G4s are defined by a positive band around 295 nm and a negative band around 260 nm. Hybrid G4 structures usually display a strong positive peak near 290 nm, often with a shoulder around 260 nm, and a negative peak near 240 nm [24, 25]. Monovalent cations, especially K^+^, are known to stabilize G4 structures by binding within their central channel [26]. The CD spectrum of the TP53-INT1 sample displayed a positive peak at 262 nm and a negative peak at 242 nm, indicating the presence of a parallel G4 structure. TP53-INT1 sample prepared without any additional KCl showed a similar spectral profile with lower peak intensities, also indicating a parallel topology (Figure 5C). The reduced CD intensity may indicate a lower extent of G4 folding since partially folded G4s can exhibit reduced peak intensities compared to fully folded G4s [27]. Compared with TP53-INT1, the mutant sequence showed markedly reduced CD signal intensity and a red shift. With and without KCl, the mutant sequence showed positive peaks at 271 and 273 nm and negative peaks at 247 and 253 nm, respectively, suggesting that the replacement of guanines with adenosines disrupted the parallel G4 structure formed by TP53-INT1.

To further examine this putative G4 structure, we performed UV-Vis thermal denaturation analysis to determine its thermal stability and melting temperature (Tm). Thermal denaturation profiles were obtained by monitoring absorbance at 295 nm with respect to temperature. G4 structures typically show hypochromism (a decrease in absorbance) at 295 nm upon heating [28]. The TP53-INT1 sequence showed hypochromism at 295 nm in both the presence and absence of additional KCl, supporting a G4 structure (Figure 5D). The Tm of the TP53-INT1 sequence was 39°C in the presence of KCl and 28°C in its absence. The higher Tm in the presence of KCl further supports the stabilizing role of K ions in the G4 structure. In contrast, the TP53-INT1 mutant sequence did not show the characteristic hypochromism associated with G4 structures. These results demonstrated that guanine-to-adenosine substitutions disrupted the G4 formation. Taken together, ThT fluorescence, CD, and UV-Vis melting profiles provided complementary evidence for the G4-forming potential of the TP53-INT1 sequence.

Having shown that the TP53-INT1 sequence can form a G4 structure *in vitro*, we next asked whether G4 stabilization in the cellular context might affect full-length transcription. We therefore treated MCF7 cells with 1 or 10 µM Pyridostatin (PDS), a selective small molecule that binds to and stabilizes G4 structures [29, 30], and assessed TP53-INT1 and full-length *TP53* mRNA levels by RT-qPCR (Figure 6). We also monitored *MYC* mRNA levels as a positive control, as stabilization of the well-characterized G4 structure in the *MYC* promoter reduces transcription [31, 32]. As expected, PDS treatment reduced *MYC* levels, consistent with promoter G4 stabilization. Interestingly, full-length *TP53* mRNA was almost completely depleted following PDS treatment, whereas TP53-INT1 levels remained largely unchanged. This differential response raised the possibility that G4 stabilization within intron 1 may have interfered with transcriptional progression beyond the G4 structure of the TP53-INT1. Of note, another G4 structure has previously been proposed within intron 3 of *TP53*, where it influences splicing patterns [33]. However, our observations suggested that G4 stabilization within the first intron may have interfered with transcription, as we did not detect full-length *TP53* transcript extending beyond the TP53-INT1 termination site, using both terminal and internal exon (exon 2-exon 3) specific primers for RT-qPCR (Figure 6).

**Figure 6.**
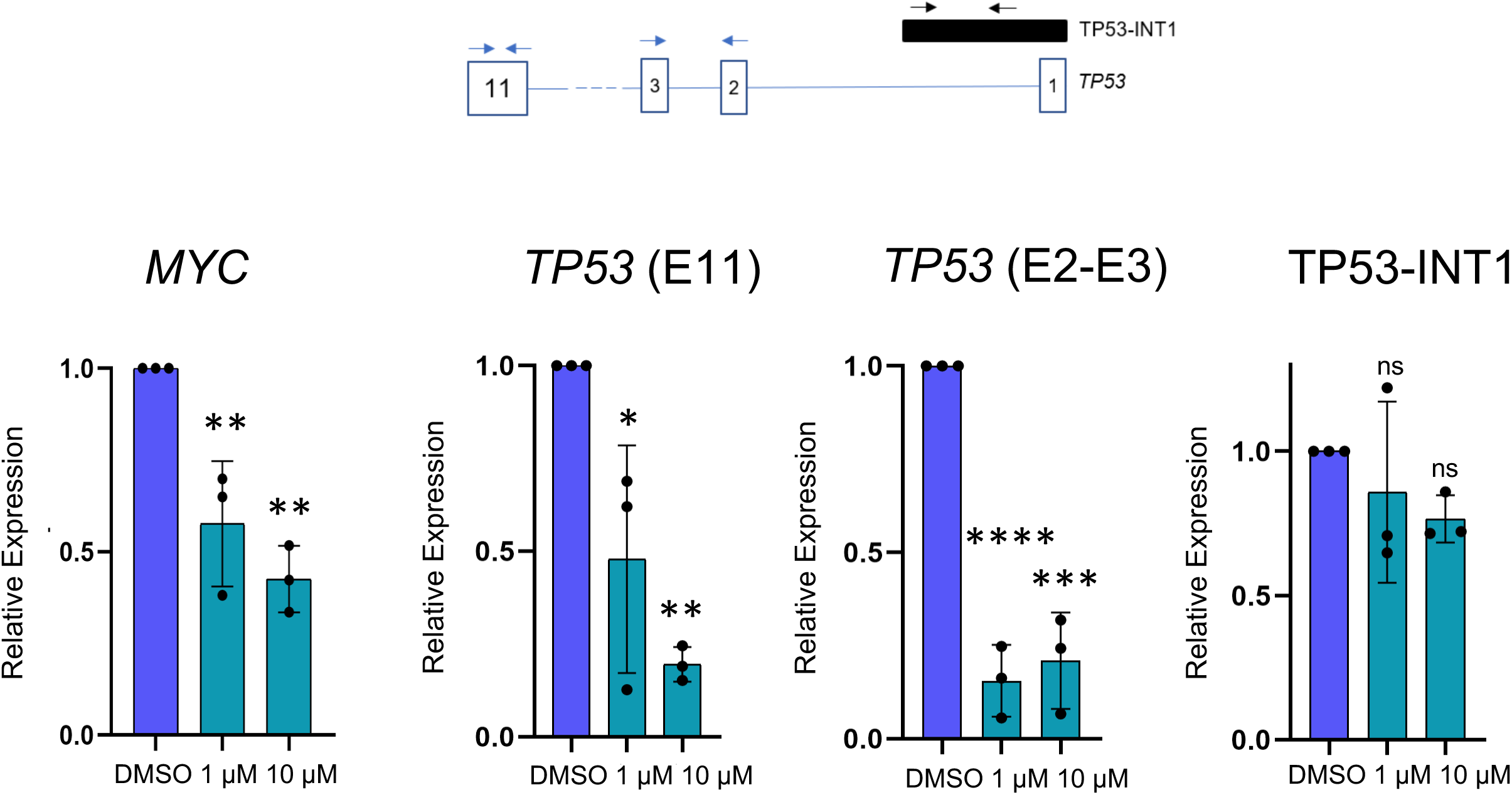
Effect of PDS stabilization of G4s. The top panel shows TP53 and TP53-INT1 gene structures and primer positions. Relative gene expression levels of *MYC*, *TP53,* and TP53-INT1 in MCF7 cells were determined after 1 µM or 10 µM PDS treatment for 24 h. *RPLP0* levels were used for normalization. Relative expression levels were then normalized to DMSO-treated samples. Data represent the mean of three independent experiments. One-way ANOVA with Tukey’s multiple comparison test was used, ns: not significant, ns: not significant, *p<0.05; **p<0.01, ***p<0.001, ****p<0.0001.

Overall, we describe a relatively stable, low-coding-potential transcript that terminates within the first intron of *TP53*. This transcript originates and terminates within an open chromatin region of intron 1 and contains a functional, canonical poly(A) signal. We also provide evidence for a potential G4 structure immediately upstream of this poly(A) site, raising the possibility that the G4 structure and an accessible poly(A) signal within TP53-INT1 may contribute to the regulation of *TP53* transcription. The function of TP53-INT1 and the regulatory mechanisms involving these intronic sequences remain to be determined.

## Discussion

First introns are often longer, more GC-rich, and more conserved than downstream introns [17, 34, 35]. They also contain a higher density of regulatory elements and functional motifs, suggesting that they can play important roles in transcriptional regulation of their host genes [17]. Here, we describe an unannotated intronic *TP53* transcript detected by DRS [6]. Long-read and DRS methods can capture full-length transcripts in individual reads, providing direct resolution of transcript structures and allowing identification of novel isoforms [36, 37] . We confirmed the 3’-end of the TP53-INT1 transcript with independent experimental and *in silico* data. Our findings are consistent with observations reported in a recent preprint, providing additional support for expression of an intronic transcript from the first intron of *TP53* [38]. We further described a functional and canonical poly(A) signal and RNA pol II occupancy patterns to support the TP53-INT1 transcript. In addition, the open chromatin architecture and a predicted G4 structure were of interest. G4 structures are formed by two or more stacked guanine tetrads, stabilized by Hoogsteen hydrogen bonds and monovalent cations such as K and Na [39, 40]. G4s are distributed across genomic regions, including promoters, exons, untranslated regions, and introns [41]. As a result, they can modulate transcription, nucleosome occupancy, chromatin accessibility, long-range chromatin interactions, and biomolecular condensate formation [42]. Hence, G4s may function as important architectural elements of chromatin throughout gene structures.

Complementing the *in vitro* characterization of the G4 structure using ThT fluorescence emission, circular dichroism and thermal denaturing profiling, treatment with pyridostatin, a G4-stabilizing drug, correlated with decreased *TP53* mRNA levels whereas TP53-INT1 levels were not affected. These results suggested that availability of a G4 structure upstream of an intronic polyadenylation signal may modulate RNA Pol II elongation and full length *TP53* transcription. Together, our findings characterize a previously unannotated *TP53* intronic transcript and reveal a local genomic context combining accessible chromatin, a functional poly(A) signal, and a potential G4-forming sequence that may influence full-length transcription. Given their length and the regulatory elements they may harbor, including poly(A) signals, characterizing noncanonical intronic transcripts could be important for understanding their potential roles in gene regulation (reviewed in [43, 44]).

Our previous study on estradiol (E2) stimulated transcriptome diversity using long read DRS [6] revealed numerous other intronically terminated transcripts. One example is a distal intronic polyadenylation of *TLE1*, assisting estrogen receptor and chromatin interactions. The resulting C-terminally truncated TLE1 protein lacked the WD-repeat domain but retained the N-terminal Q-domain required for dimerization and tetramerization, illustrating how alternative transcript termination can alter protein domain composition without eliminating protein interaction capabilities. Together, these findings support the notion that the functional consequences of intronic polyadenylation may depend on the position of the poly(A) site within a gene and the structural features of the resulting transcript or protein isoform. Intronic polyadenylation may therefore contribute to gene regulation, highlighting the importance of characterizing intronic transcripts in their genomic and molecular context. In the context of the first intron of *TP53*, we provide evidence on mechanisms leading to the expression of TP53-INT1 transcript but the biological function of TP53-INT1 itself and formation, stability, or potential regulatory activity of the G4 structure identified in this region remain to be further investigated.

## Materials and Methods

### Cell Culture and Treatments

Breast cancer cell lines were cultured in Dulbecco’s Modified Eagle Medium (DMEM) supplemented with 10% Fetal Bovine Serum (FBS) (Biowest, S1810-500), 2% L-Glutamine, 1% Sodium Pyruvate Solution and 1% Penicillin/Streptomycin (BI, 03-031-1B). Cells were incubated at 37 °C with 95% humidified air and 5% CO_2_ and were grown as monolayers. Normal breast tissue cDNA was used from the Breast Cancer cDNA Array IV (Origene). TurboFect Transfection Reagent (Thermo Fisher Scientific) was used for transfections. Pyridostatin (Sigma-Aldrich SML 2690) was solubilized in DMSO.

### mRNA Half-life

Total RNA was isolated using the High Pure RNA Isolation Kit (Roche Applied Science) and treated with DNase I (Roche Applied Science). cDNA was synthesized using the RevertAid First Strand cDNA Synthesis Kit (Thermo Fisher Scientific). To determine mRNA half-lives, cells were treated with actinomycin D (10 µg/mL, Tocris Bioscience) to inhibit transcription, and RNA levels were subsequently monitored over time.

### RT-qPCR and 3’RACE

RNA isolation, quantification, and cDNA synthesis were conducted according to the MIQE guidelines [45]. RNA samples were DNase-treated (Thermo Fisher Scientific, EN0521) to eliminate DNA contamination. Lack of DNA contamination was controlled and verified by PCR using RNA samples as template. SsoAdvanced Universal SYBR Green Supermix (BioRad) and CFX Connect Real-Time PCR were used. Fold changes were normalized to the *RPLP0* reference gene. For relative quantification, the reaction efficiency was calculated using the ΔΔCt formula [34]. To ensure reaction specificity, the melt curve of each reaction was analyzed. Rapid amplification of cDNA ends (3’RACE)-specific cDNA synthesis was performed as described [46]. For site-directed mutagenesis, TP53-INT1 poly(A) motif (6 bp) was deleted using primers designed with QuikChange Primer Design tool and as described [46]. All primer sequences are given in the Supplementary File (Table S1).

### Dual Luciferase Assay for Poly(A) Activity

*Firefly* luciferase reporter system (pMIR-Report, pMIR) and pMIR-ΔpA which lacks the SV40 poly(A) signal were used to test poly(A) signals. PCR cloned poly(A) signal regions were confirmed by sequencing (pMIR Forward: 5’-AGGCGATTAAGTTGGGTA-3’, Reverse: 5’-GGAAAGTCCAAATTGCTC-3’). MCF7 cells were co-transfected with 25 ng phRL-TK (Renilla Luciferase) and 225 ng pMIR or pMIR Δpoly(A) (*Firefly* Luciferase) vectors. 50,000 cells were seeded 24 hours before transfection using TurboFect Transfection Reagent (Thermo Fisher). After 24 hours, the Dual-Luciferase Reporter Assay (Promega) protocol was followed, and luminescence was measured using the Modulus Microplate Luminometer (Turner Biosystems). *Firefly/Renilla* ratios were calculated and normalized to those of cells transfected with the empty pMIR vector.

### In silico Analysis

Coding Potential Calculator (CPC2) was used to evaluate the coding potential of TP53-INT1 [8]. G4 Hunter (https://bioinformatics.ibp.cz/#/) and G4Catchall (homes.izmirekonomi.edu.tr/odoluca/G4Catchall/) were used to predict G4 structures [18, 19]. Cistrome Data Browser (http://cistrome.org/) was used to investigate RNA pol II occupancy and open chromatin histone marks [16]. Integrative Genomics Viewer (IGV) was used for visualization. RNA sequencing data for the Genotype Tissue Expression project (GTEx; https://gtexportal.org/home/) was used to report *TP53* mRNA isoforms in mammary tissue.

### G4 Testing

Oligonucleotides of the predicted G-quadruplex (G4) forming sequence (TP53-INT1), its complementary strand (TP53-INT1comp), mutant sequence (TP53-INT1-mut), Pu27-G4 forming sequence of the MYC promoter [22], and dA_32_) were purchased from Oligomer Biotechnology Inc. (Ankara, Türkiye). Concentrations of oligonucleotide stock solutions were determined according to the Beer-Lambert law using a JASCO V-730 UV–Vis spectrophotometer by measuring the absorbance at 260 nm. The molar extinction coefficient values given in the Table S2 were obtained using IDT’s OligoAnalyser Tool. All nucleic acid samples were prepared in 25 mM K-phosphate buffer at pH 7.0 in the presence of 70 mM KCl in Millipore water, where the DNA concentration was 3.0 µM per strand. Thioflavin T molecule (ThT) was purchased from Sigma Aldrich (St. Louis, MO, USA), and stock solution was prepared in Millipore water. Concentration was determined by UV-Vis spectroscopy using a molar extinction coefficient of 36,000 M□¹ cm□¹ [47]. For the fluorescence experiments, 1.0 µM ThT and 3.0 µM oligos were used [48].

### Fluorescence Emission

Fluorescence spectroscopy was used to evaluate G4 formation using ThT, which serves as a selective G4 probe due to its pronounced fluorescence enhancement upon binding to G4 structures over other DNA secondary conformations [20]. Fluorescence emission spectra were collected using a Cary Eclipse fluorescence spectrophotometer with 10 mm quartz cuvettes. Using an excitation wavelength of 412 nm for ThT, emission spectra were recorded over the range of 430–700 nm, with excitation and emission slit widths of 5.0 nm, a photomultiplier tube voltage of 700 V, and a scan rate of 600 nm/min.

### Circular Dichroism (CD) Spectrophotometry

Circular dichroism (CD) spectroscopy was used to assess G4 formation and determine the overall topology of the secondary structures [27]. CD spectra were recorded between 200 nm and 500 nm at 15°C using a JASCO J-1500 CD spectropolarimeter equipped with a Peltier temperature control unit.

### Thermal Denaturation Profiling

Thermal denaturation experiments were performed to verify the formation of stable G4 structures and determine their stability. First, oligonucleotides samples prepared were heated at 93°-95°C for 5 min and slowly cooled to room temperature overnight to allow proper secondary structure formation. Next, thermal denaturation profiles were obtained by monitoring changes in UV-Vis absorbance at 295 nm while increasing the sample temperature from 15°C to 95°C at a rate of 2°C per minute [49]. Agilent Cary 8454 UV-Vis spectrophotometer equipped with an Agilent 89090A Peltier temperature control unit was used.

## Author Contributions

AEEB conceived the study and supervised experiments; OPC supervised experiments, ICE, IG and EAY performed experiments and analyzed data. All authors contributed to manuscript preparation.

## Acknowledgement

This project was funded by TÜSEB TB-37875 and TÜBİTAK 122Z679. EAY was supported by TUBITAK BIDEB 2211A. We would like to thank Kübra Demirbilek for her assistance in the initial characterization of the G4 structures and Prof. Dr. Mesut Muyan for insightful discussions.

