## Supplementary material for "A G-Quadruplex Containing First Intron Harbors a Functional Polyadenylation Signal in *TP53*": Figure S

### Supplementary Figures

## A

>546fd2ff-dcfd-4f49-be47-06cfb26023e0_17:7588000

CTAGAGCCACCGTCCAGGGAGCAGGTAGCTGCTGGGCTCCGGGGACACTTTGCGTTCGGGCTGGGAGCGTGCTTTCCACGACGGTGACACGCTTCCCTGGATTGGGTAAGCTCC
TGACTGAACTTGATGAGTCCTCTCTGAGTCACGGGCTCTCGGCTCCGTGTATTTTCAGCTCGGGAAAATCGCTGGGGCTGGGGGTGGGGCAGTGGGGACTTAGCGAGTTTGGGG
GTGAGTGGGATGGAAGCTTGGCTAGAGGGATCATCATAGGAGTTGCATTGTTGGGAGACCTGGGTGTAGATGATGGGGATGTTAGGACCATCCGAACTCAAAGTTGAACGCCT
AGGCAGAGGAGTGGAGCTTTGGGGAACCTTGAGCCGGCCTAAAGCGTACTTCTTTGCACATCCACCCGGTGCTGGGCGTAGGGAATCCCTGAAATAAAAGATGCACAAAGCAT
TGAGGTCTGAGACTTTTGGATCTCGAAACATTGAGAACTCATAGCTGTATATTTTAGAGCCCATGGCATCCTAGTGAAAACTGGGGCTCCATTCCGAAATGATCATTTGGGGGT
GATCCGGGGAGCCCAAGCTGCTAAGGTCCCACAACTTCCGGACCTTTGTCCTTCCTGGAGCGATCTTTCCAGGCAGCCCCCGGCTCCGCTAGATGGAGAAAATCCAATTGAAGG
CTGTCAGTCGTGGAAGTGAGAAGTGCTAAACCAGGGGTTTGCCCGCCAGGCCGAGGAGGACCGTCGCAATCTGAGAGGCCCGGCAGCCCTGTTATTGTTTGGCTCCACATTTAC
ATTTCTGCCTCTTGCAGCAGCATTTCCGGTTTCTTTTTGCCGGAGCAGCTCACTATTCACCCGATGAGAGGGGAGGAGAGAGAGAGAAAATGTCCTTTAGGCCGGTTCCTCTTAC
TTGGCAGAGGGAGGCTGCTATTCTCCGCCTGCATTTCTTTTTCTGGATTACTTAGTTATGGCCTTTGCAAAGGCAGGGGTATTTGTTTTGATGCAAACCTCAATCCCTCCCCTTCT
TTGAATGGTGTGCCCCACCCCGCGGGTCGCCTGCAACCTAGGCGGACGCTACCATGGCGTGAGACAGGGAGGGAAAGAAGTGTGCAGAAGGCAAGCCCGGAGGTATTTTCAA
GAATGAGTATATCTCATCTTCCCGGAGGAAAAAAAAAAAAGAATGGGTACGTCTGAGAATCAAATTTTGAAAGAGTGCAATGATGGGTCGTTTGATAATTTGTCGGAAAAACAA
TCTACCTGTTATCTAGCTTTGGGCTAGGCCATTCCAGTTCCAGACGCAGGCTGAACGTCGTGAAGCGGAAGGGGCGGGCCCGCAGGCGTCCGTGTGGTCCTCCGTGCAGCCCTC
CGGCCCGAGCCGGTTCTTCCTGGTAGGAGGCGGAACTCGAATTCATTTCTCCCGCTGCCCCATCTCTTAGCTCGCGGTTGTTTCATTCCGCAGTTTCTTCCCATGCACCTGCCGCG
TACCGGCCACTTTGTGCCGTACTTACGTCATCTTTTTCCTAAATCGAGGTGGCATTTACACACAGCGCCAGTGCACACAGCAAGTGCACAGGAAGATGAGTTTTGGCCCCTAACC
GCTCCGTGATGCCTACCAAGTCACAGACCCTTTTCATCGTCCCAGAAACGTTTCATCACGTCTCTTCCCAGTCGATTCCCGACCCCACCTTTATTTTGATCTCCATAACCATTTTG
CCTGTTGGAGAACTTCATATAGAATGGAATCAGGCTGGGCGCTGTGGCTCACGCCTGCACTTTGGGAGGCCGAGGCGGGCGGATTACTTGAGGATAGGAGTTCCAGACCAGCG
TGGCCAACGTGGTGAATCCCCGTCTCTACTAAAAAATACAAAAATTAGCTGGGCGTGGTGGGTGCCTGTAATCCCAGCTATT*CGGGAGGGTGAGGCAGGAGAA*TCGCTTGAACC
CGGGAGGCAGAGGTTGCAGTGAGCCAAGATCGTGCCACTACACTCCAGCCTGGGCGACAAGAACGAAACTCCGTCTCAAAAAAAAGGGGGGAATCATACATTATGTGCTCATT
TTTGTCGGGCTTCTGTCCTTCAATGTACTGTCTGACATTCGTTCATGTTGTATATATCAGTATTTTGCTCCTTTTCATTTAGTATAGTCCATCGATTGTATATCCGTCCTTTTGATGG
CCTTTTGAGTTGTTTCCCATTTGCGGTTATGAAATAAAGCTGCTATAAACATTCTTGTACAA

## B

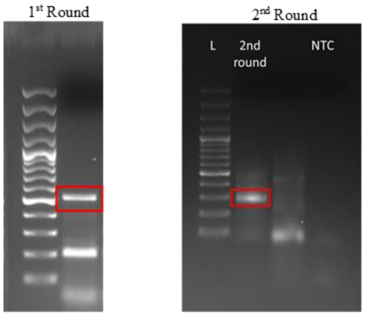


## C

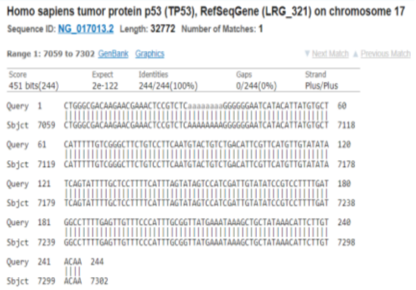


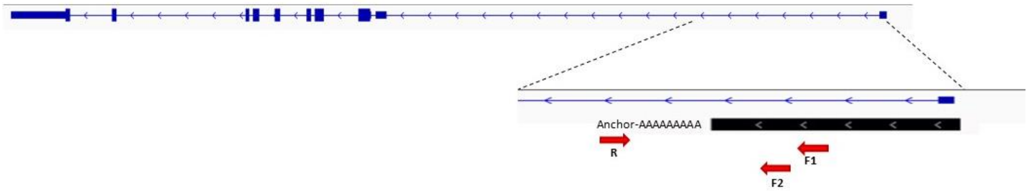


## D

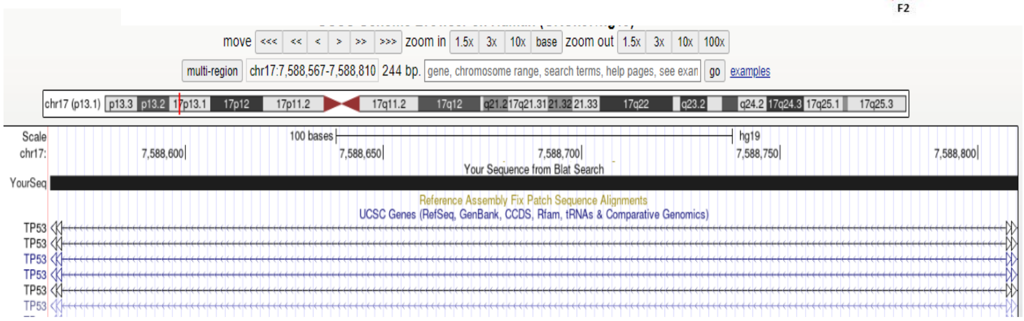


**Figure S1. TP53-INT1 A.** Sequence of the ONT DRS identified transcript, **B.** Confirmation by 3'RACE. First and second round 3'RACE PCR result, **C.** Boxed PCR band was purified and cloned into pGEM-T vector and sequenced, **D**. Cloned 3' RACE product (black) in alignment with *TP53* intron 1

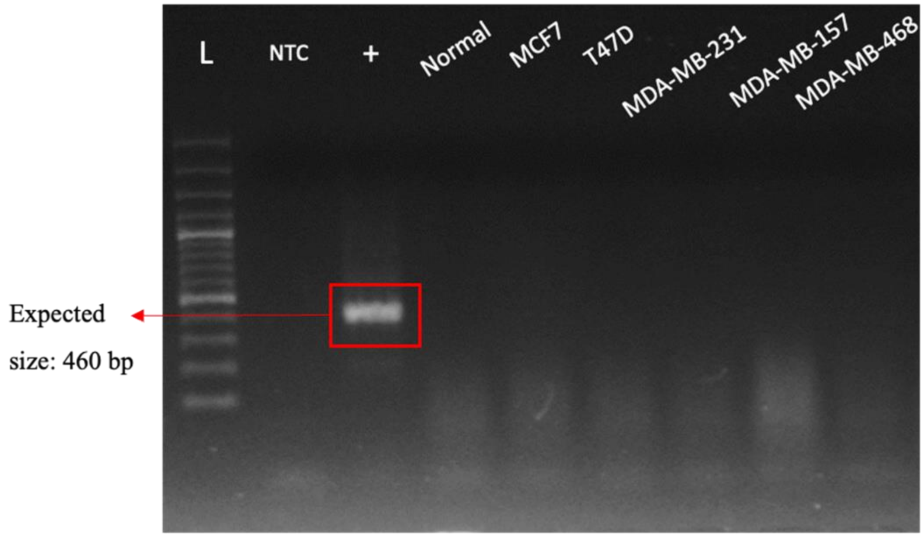


**Figure S2. Representative example for the absence of DNA contamination in RNA samples.** Lack of DNA contamination was confirmed by using RNA samples as templates for PCR using *GAPDH* primers. NTC (no template control), Lane L represents the DNA Ladder, Lane + is positive control (gDNA template), the remaining lanes represent RNA samples that were treated with DNase.

**A** **TP53-INT1**

| ID | Label | Coding probability | Peptide length(aa) | Fickett score | Isoelectric point | ORF integrity | Details |
|---|---|---|---|---|---|---|---|
| 546fd2ff-dcfd-4f49-be47-06cfb26023e0_17:7588000 | noncoding | 0.25598 | 140 | 0.29707 | 9.91741943359 | complete | View |

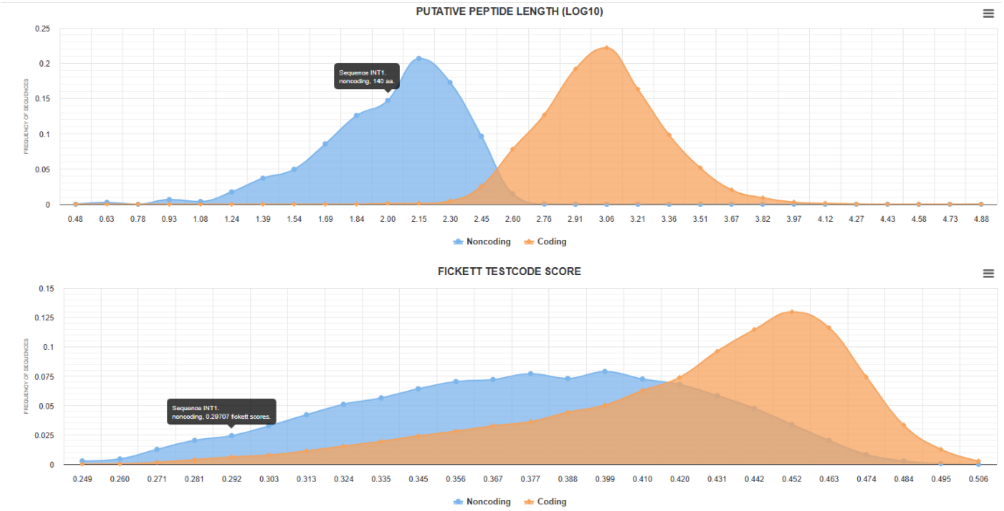


**B** **TP53**

| ID | Label | Coding probability | Peptide length(aa) | Fickett score | Isoelectric point | ORF integrity | Details |
|---|---|---|---|---|---|---|---|
| TP53 | coding | 1 | 394 | 0.35661 | 6.33172607422 | complete | View |

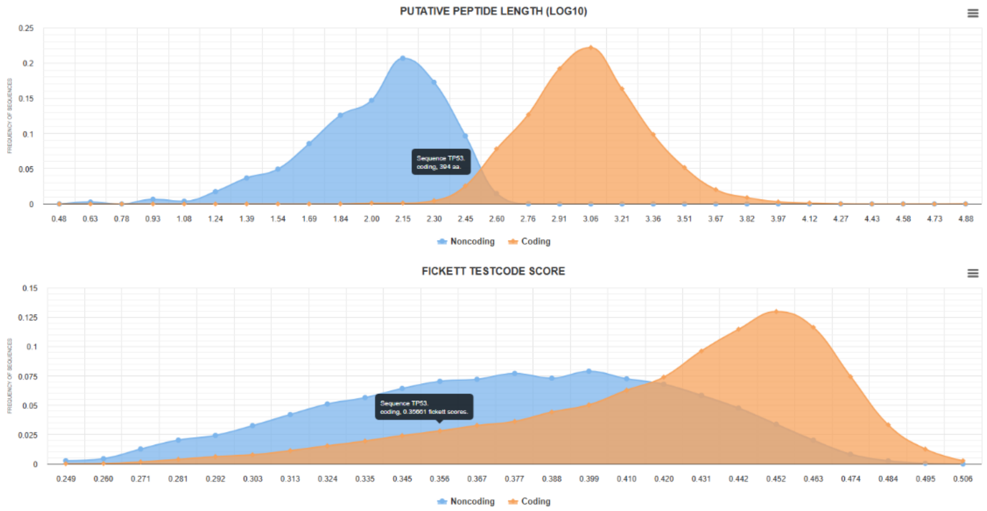


**Figure S3.** Coding Potential Calculator (CPC2) analysis of **A**.TP53-INT1 and **B**. *TP53* mRNA. Summary tabular output and graphical view of features' distributions are shown.

**A**

| | |
|---|---|
| TP53 | 5' - …CATTTTTATATCGATCTCTTATTTTAC**<u>AATAAA</u>**ACTTTGCTGCCACCTGTGTGTCTGAGG… - 3' |
| TP53-INT1 | 5' - …TCCCATTTGCGGTTATGA**<u>AATAAA</u>**GCTGC**TATAAA**CATTCTTGTACAATTCTTTTT… - 3' |
| TP53-INT1 ΔpA | 5' - …TCCCATTTGCGGTTATGA**------------**GCTGC**TATAAA**CATTCTTGTACAATTCTTTTT… - 3' |

**B**

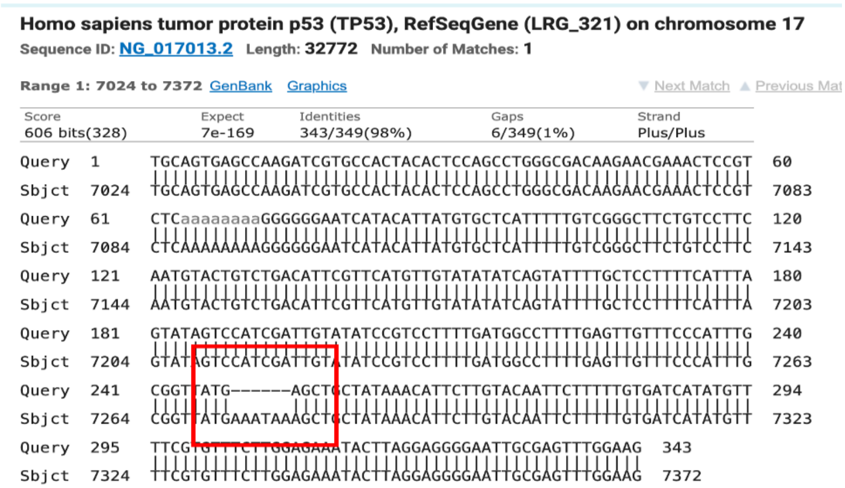


**Figure S4. A.** poly(A) signals in the terminal exon of *TP53*, and TP53-INT1 are shown. TP53-INT1 ΔpA was generated by site directed mutagenesis to delete the canonical AATAAA sequence, **B**. Sequencing and blast confirms the deletion of the AATAAA signal in the TP53-INT1- ΔpA construct.

**A**

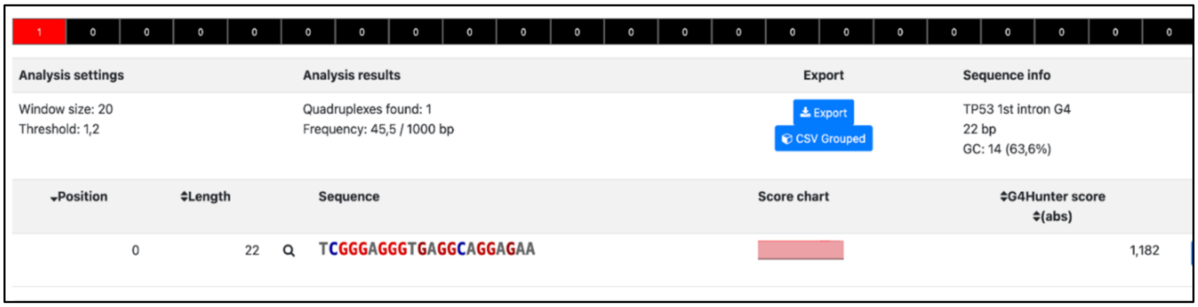


**B**

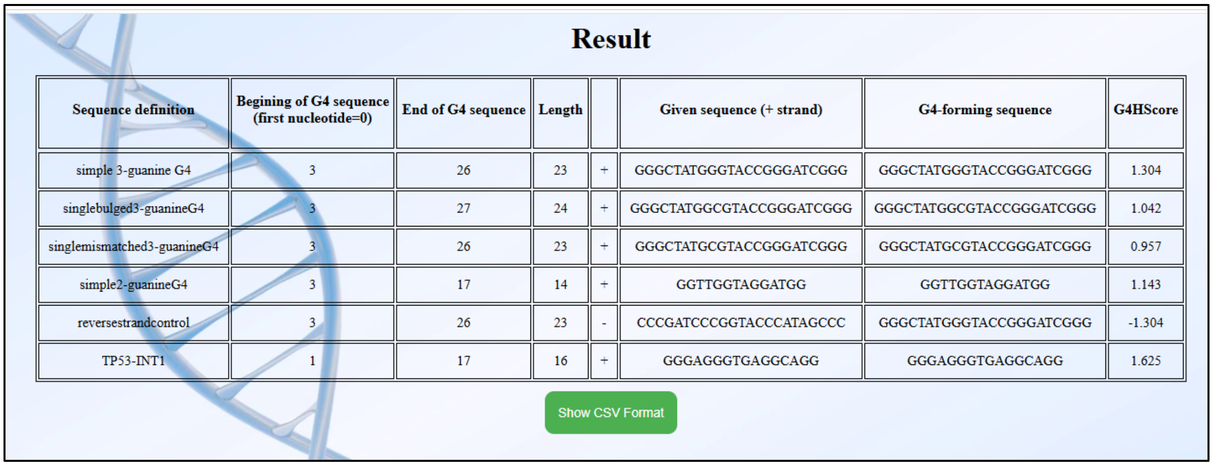

Result

| Sequence definition | Begining of G4 sequence (first nucleotide=0) | End of G4 sequence | Length | | Given sequence (+ strand) | G4-forming sequence | G4HScore |
|---|---|---|---|---|---|---|---|
| simple 3-guanine G4 | 3 | 26 | 23 | + | GGGCTATGGGTACCGGGATCGGG | GGGCTATGGGTACCGGGATCGGG | 1.304 |
| singlebulged3-guanineG4 | 3 | 27 | 24 | + | GGGCTATGGCGTACCGGGATCGGG | GGGCTATGGCGTACCGGGATCGGG | 1.042 |
| singlemismatched3-guanineG4 | 3 | 26 | 23 | + | GGGCTATGCGTACCGGGATCGGG | GGGCTATGCGTACCGGGATCGGG | 0.957 |
| simple2-guanineG4 | 3 | 17 | 14 | + | GGTTGGTAGGATGG | GGTTGGTAGGATGG | 1.143 |
| reversestrandcontrol | 3 | 26 | 23 | - | CCCGATCCCGGTACCCATAGCCC | GGGCTATGGGTACCGGGATCGGG | -1.304 |
| TP53-INT1 | 1 | 17 | 16 | + | GGGAGGGTGAGGCAGG | GGGAGGGTGAGGCAGG | 1.625 |

Show CSV Format

**Figure S5.** *In silico* predictions by **A**. G4Hunter, **B**. G4Catchall (top 5 sequences were control sequences inserted by the tool).

#### Table S1. Primer sequences

| **Gene** | **Forward (5'-3')** | **Reverse (5'-3')** |
|---|---|---|
| *TP53-INT1* | ATTCCAGTTCCAGACGCAGG | AACCGCGAGCTAAGAGATGG |
| *TP53* | TCCCCTGCCATTTTGGGTTT | GCAGGCCAACTTGTTCAGTG |
| TP53-INT1 3'RACE (1) | CCATTTTGCCTGTTGGAGAACTT | GACCACGCGTATCGATGTCGACTTTTTTTTTTTTTTTT |
| TP53-INT1 3'RACE (2) | CTGGGCGACAAGAACGAAAC | GACCACGCGTATCGATGTCGACTTTTTTTTTTTTTTTT |
| *TP53-INT1 pA* Cloning | CGCATGAGCTCTATCCGTCCTTTTGATGGCC | CGCATACGCGTCTTCCAAACTCGCAATTCCC |
| *TP53-pA* Cloning | CGCATGAGCTCCATTCTGCAAGCACATCTGC | CGCATACGCGTCACACTCATTGCAGACTCAG |

#### Table S2. Oligonucleotides and extinction coefficients

| Oligonucleotide Name | Sequence 5'-3' | Molar Extinction Coefficient (L/(mol.cm)) |
|---|---|---|
| TP53-INT1 | CGGGAGGGTGAGGCAGGAGAA | 231300 |
| TP53-INT1 comp | GCCCTCCCACTCCGTCCTCTTA | 183500 |
| TP53-INT1 mutant | CGAGAGAATGAGACAAGAGAA | 242600 |
| $dA_{32}$ | AAAAAAAAAAAAAAAAAAAAAAAAAAAAAAAA | 387400 |
| MYC | TGGGGAGGGTGGGGAGGGTGGGGAAGG | 279900 |
